# Multivariate genotypic analyses that identify specific genotypes to characterize disease and control groups in ADNI

**DOI:** 10.1101/235945

**Authors:** Derek Beaton, Jenny R. Rieck, Fahd Alhazmi, ADNI, Hervé Abdi

## Abstract

**INTRODUCTION:** Genetic contributions to Alzheimer’s Disease (AD) are likely polygenic and not necessarily explained by uniformly applied linear and additive effects. In order to better understand the genetics of AD, we require statistical techniques to address both polygenic and possible non-additive effects.

**METHODS:** We used partial least squares-correspondence analysis (PLS-CA)—a method designed to detect multivariate genotypic effects. We used ADNI-1 (*N* = 756) as a discovery sample with two forms of PLS-CA: diagnosis-based and ApoE-based. We used ADNI-2 (*N*= 791) as a validation sample with a diagnosis-based PLS-CA.

**RESULTS:** With PLS-CA we identified some expected genotypic effects (e.g., *APOE/TOMM40*, and *APP)* and a number of new effects that include, for examples, risk-associated genotypes in *RBFOX1* and *GPC6* and control-associated genotypes in *PTPN14* and *CPNE5*.

**DISCUSSION:** Through the use of PLS-CA, we were able to detect complex (multivariate, genotypic) genetic contributions to AD, which included many non-additive and non-linear risk and possibly protective effects.

## 1. Introduction

Many genes have been linked to Alzheimer’s disease (AD) such as SORL1 [1], EPHA1 [2], CLU, and PICALM [3], all of which have shown effects in a variety of studies [4]. Though there are well-known genetic effects in AD [5–7]—there are still some controversial findings. For example, the work reported in [8] showed effects of rare variants in PLD3, where follow up studies showed both replication [9] and no replication [10] of PLD3 effects. Overall, recent works suggest that the genetic contributions to AD are: polygenic [11,12], epistatic [13], and non-linear or non-additive [14]. However, the routine analytical approaches for genetic and genomic data do not accommodate such complexities; but, if the genetics of AD are so complex why are we still using statistical methods whose assumptions are not suited to detect such complex effects?

With the advent of genome-wide studies, single nucleotide polymorphisms (SNPs) are almost exclusively analyzed with the additive model. The additive model transforms a SNP from base pair letters into a count based on number of minor alleles where usually a major homozygote is “0,” a heterozygote is “1,” and a minor homozygote is “2.” The additive model has become popular because it is viewed as both a practical approach [15] and a suitable model for complex traits [16]. However, the assumptions of additivity and/or linearity do not always hold (see [17,18]). Recent work (in cholesterol) has shown that the genotypic (“full”) model is better than the additive model to detect genetic contributions to complex traits [19]. Furthermore, additive assumptions can be detrimental because the *a priori* choice of a model that does not match the true inheritance pattern causes a loss of power [20]. Additionally, the values of ‘0’ and ‘2’ are inherently ambiguous across samples, and this ambiguity could lead to misinterpretation or even the dismissal of effects because of the direction of genotypic effects. For possible examples in AD: in [21] the authors reported “direction changes” in their own replication analyses, whereas in [22] the authors report effects in the opposite direction in their attempt to replicate work in [8]. The additive model has been used in many AD studies even though the existence of non-additive effects is often acknowledged and expected [23,24]. With resources such as the Alzheimer’s Disease Neuroimaging Initiative (ADNI), researchers have had the opportunity to analyze genome-wide data in a variety of ways, yet the ADNI genome-wide data have almost exclusively been analyzed with the additive model (see, e.g., [25–30]).

In this study we investigated the ADNI genome-wide data with a technique tailored for complex and polygenic effects called partial least squares-correspondence analysis (PLS-CA, [31]). PLS-CA is a multivariate technique that allows for a more general approach (i.e., genotypic model). Furthermore, PLS-CA was designed to address the complexity of genetic contributions (i.e., polygenicity, non-additivity) so that we can detect, rather than assume, specific genotypic effects. PLS-CA treats SNPs as categorical data where the genotpyes are levels within those SNP categories. Our study was designed to identify multiple specific genotypic effects for AD (possible risk factors) and for controls (possible protective factors); the specificity afforded by PLS-CA helps disentangle some of the complexities of genetic contributions to AD. Our goals with this study were two-fold: (1) apply the technique within AD in order to potentially reveal some of the genetic complexities of AD and (2) illustrate a novel approach to perform genetic and genomic association studies.

Our paper is outlined as follows. In *Methods*, we describe the ADNI data used in this study, followed by descriptions of the SNP and genotype quality control, and statistical techniques. We then detail the two phases of our study: (1) “Discovery” (with ADNI-1) which includes two genome-wide association analyses (one based on diagnosis, and another based on *APOE)*, and (2) “Validation” (with ADNI-GO/2) that used the genotypes identified in “Discovery” to create a candidate panel and then test group association in a new sample. In *Results* we present findings from each of the phases. In *Discussion* we emphasize the “Validation” phase followed by conclusions and limitations.

## 2. Methods

Data used in the preparation of this article come from the ADNI database (http://adni.loni.usc.edu). The ADNI project was launched in 2003 as a public-private funding partnership and includes public funding by the National Institute on Aging, the National Institute of Biomedical Imaging and Bioengineering, and the Food and Drug Administration. The primary goal of ADNI has been to collect a wide variety of measures to assess the progression of mild cognitive impairment (MCI) and early Alzheimer’s disease (AD). ADNI is the result of efforts of many co-investigators from a broad range of academic institutions and private corporations. Michael W. Weiner, MD (VA Medical Center and University of California at San Francisco) is the ADNI Principal Investigator. Subjects have been recruited from over 50 sites across the U.S. and Canada. For up-to-date information, see www.adni-info.org.

Our study included genomic, APOE, and diagnostic data from ADNI-1 and ADNI-GO/2 in the AD, MCI, and control (CON) groups. ADNI-1 used the Illumina Human610-Quad BeadChip while ADNI-GO/2 used the Illumina HumanOmniExpress BeadChip.

### 2.1 Participants

We obtained final totals of 756 participants from ADNI-1 (AD = 344, MCI = 204, CON = 208) and 791 participants from ADNI-GO/2 (AD = 203, MCI = 319, CON = 269). Table 1 includes overviews (e.g., demographics, cognitive measures) of the ADNI-1 and ADNI-GO/2 cohorts. Demographic, cognitive, and diagnostic measures were retrieved from the ADNIMERGE package (available via http://adni.loni.usc.edu/). Table 1 generally includes measures collected at baseline, though for this study we used the last available diagnosis. Not all measures were available at all time points, thus we characterize the sample by the baseline data and last available diagnosis. ADNI-GO/2 had additional recruitment groups: subjective memory complaints (SMC) and MCI was split into early and late. For the ADNI-GO/2 diagnoses in our analyses the two MCI groups were combined. The SMC category was not used as a diagnosis in later time points in the ADNI-GO/2 study, only as a recruitment group at baseline. See Table 1 for details.

**Table 1.** Overview of ADNI 1 and ADNI-GO/2 cohorts.

|  | *Total* | CON-b | MCI-b | AD-b |
| --- | --- | --- | --- | --- |
| *Control* | 208 | 195 | 13 | 0 |
| *Mild Cognitive Impairment* | 204 | 15 | 188 | 1 |
| *Alzheimer's Disease* | 344 | 3 | 163 | 178 |

(b) Demographics and summaries for ADNI1
|  | Diagnosis<br>(Female) | Age-b mean<br>(s.d.) | Education-b mean<br>(s.d.) | MMSE-b mean<br>(s.d.) | CDRSB-b mean<br>(s.d.) |
| --- | --- | --- | --- | --- | --- |
| *Control* | 208 (93) | 75.36 (5.27) | 16.07 (2.77) | 29.01 (1.14) | 0.10 (0.33) |
| *Mild Cognitive Impairment* | 204 (70) | 74.89 (7.40) | 15.61 (3.18) | 27.41 (1.80) | 1.34 (0.83) |
| *Alzheimer's Disease* | 344 (145) | 75.22 (7.12) | 15.19 (3.09) | 24.96 (2.57) | 3.11 (1.87) |

(c) Diagnosis changes for ADNI-GO/2
|  | *Total* | CON-b | SMC-b | eMCI-b | IMCI-b | AD-b |
| --- | --- | --- | --- | --- | --- | --- |
| *Control* | 269 | 142 | 96 | 25 | 6 | 0 |
| *Mild Cognitive Impairment* | 319 | 12 | 3 | 227 | 76 | 1 |
| *Alzheimer's Disease* | 203 | 1 | 0 | 25 | 52 | 125 |

(d) Demographics and summaries for ADNI-GO/2
|  | Diagnosis<br>(Female) | Age-b mean<br>(s.d.) | Education-b mean<br>(s.d.) | MMSE-b mean<br>(s.d.) | CDRSB-b mean<br>(s.d.) |
| --- | --- | --- | --- | --- | --- |
| *Control* | 269 (147) | 72.37 (6.02) | 16.68 (2.50) | 29.07 (1.19) | 0.17 (0.45) |
| *Mild Cognitive Impairment* | 319 (138) | 71.93 (7.77) | 16.06 (2.64) | 28.17 (1.68) | 1.23 (0.78) |
| *Alzheimer's Disease* | 203 (82) | 73.79 (7.88) | 15.83 (2.63) | 24.76 (2.92) | 3.67 (1.86) |

(e) Self-identified race (non-Hispanic/Latino) for ADNI1 and ADNI-GO/2
|  | <u>ADNI1</u> | <u>ADNI-GO/2</u> |
| --- | --- | --- |
| *White* | 703 (685) | 732 (700) |
| *Asian* | 12 (12) | 12 (12) |
| *Black* | 37 (36) | 30 (30) |
| *American Indian/Alaskan* | 1 (1) | 2 (2) |
| *Hawaiian/Other Pacific Islander* | 0 (0) | 2 (2) |
| *Unknown/More than 1* | 3 (1) | 13 (7) |
| **Total** | **756 (735)** | **791 (753)** |

*Note.* Overviews of diagnostic, demographic, and cognitive/behavioral information for ADNI 1 and ADNI-GO/2. All information retrieved from the ADNIMERGE package. CON = control, SMC = subjective memory complaints, MCI = mild cognitive impairment, eMCI = early MCI, IMCI = late MCI, AD = Alzheimer's Disease, MMSE = Mini-mental state exam, CDRSB = Clinical Dementia Rating-Sum of Boxes. Items denoted with '-b' are baseline measures. (a) shows last available diagnosis vs. diagnosis at baseline for the ADNI-1 cohort. (b) shows the demographic and cognitive/behavioral measures for ADNI1. (b) shows last available diagnosis vs. diagnosis at baseline for the ADNI-GO/2 cohort. (d) shows the demographic and cognitive/behavioral measures for ADNI-GO/2. Note that ADNI-GO/2 included two subtypes of MCI (early and late) as well as the SMC category. (e) shows self-identified race and ethnicity for both ADNI1 and ADNI-GO/2.

### 2.2 Statistical techniques

Preprocessing, analyses, and graphics were performed primarily in R [32] with the `Exposition, TExPosition`, and `TInPosition` packages [33,34]. Some in-house MATLAB (Mathworks Inc., Natick, MA) code was used for resampling. Code available in Github (R packages: https://github.com/derekbeaton/ExPosition-Family/; `MATLAB` code: https://github.com/derekbeaton/Misc).

Because SNPs are categorical we required particular multivariate techniques designed specifically for categorical data. The primary techniques used in this study—multiple correspondence analysis (MCA) and PLS-CA—are analogous to principal components analysis (PCA) and partial least square correlation (PLSC) but are designed to handle categorical data and generally operate under the assumptions of χ^2^. Data were recoded from nominal (categorical) to disjunctive format (see Table 2) because with this format PLS-CA can analyze data with the genotypic model under the assumptions of χ^2^. We used (MCA) instead of PCA to correct for stratification effects. MCA is the analog of PCA—that is, a method that produces orthogonal, rank-ordered by variance components—but designed for data in a disjunctive format and also adheres to the assumptions of χ^2^.

**Table 2.** Nominal and disjunctive formats of data.

|  | SNP1 (with minor homozygote > 5%) | SNP2 (with minor homozygote < 5%) |
| --- | --- | --- |
| *Subj.1* | AG | CA |
| *Subj.2* | AA | CA |
| ... | ... | ... |
| *Subj.i* | AG | CC |
| ... | ... | ... |
| *Subj.I-1* | <NA> | AA |
| *Subj.I* | GG | AA |

(b) Disjunctive SNPs
|  | SNP1 (minor homozygote > 5%) |  |  | SNP2 (minor homozygote < 5%) |  |
| --- | --- | --- | --- | --- | --- |
|  | AG | AA | GG | CA+CC | AA |
| *Subj.1* | 1 | 0 | 0 | 1 | 0 |
| *Subj.2* | 0 | 1 | 0 | 1 | 0 |
| ... | ... | ... | ... | ... | ... |
| *Subj.i* | 1 | 0 | 0 | 0 | 0 |
| ... | ... | ... | ... | ... | ... |
| *Subj.I-1* | .2 | .7 | .1 | 0 | 1 |
| *Subj.I* | 0 | 0 | 1 | 0 | 1 |

(c) Dx and ApoE
|  | Dx | #ApoE E4 |
| --- | --- | --- |
| *Subj.1* | AD | 2 |
| *Subj.2* | AD | 1 |
| ... | ... | ... |
| *Subj.i* | MCI | 1 |
| ... | ... | ... |
| *Subj.I-1* | MCI | 2 |
| *Subj.I* | CN | 0 |

(d) Disjunctive Dx and ApoE
|  | Dx |  |  | # APOE-E4 Alleles |  |  |
| --- | --- | --- | --- | --- | --- | --- |
|  | AD | MCI | CN | 2 | 1 | 0 |
| *Subj.1* | 1 | 0 | 0 | 1 | 0 | 0 |
| *Subj.2* | 1 | 0 | 0 | 0 | 1 | 0 |
| ... | ... | ... | ... | ... | ... | ... |
| *Subj.i* | 0 | 1 | 0 | 0 | 1 | 0 |
| ... | ... | ... | ... | ... | ... | ... |
| *Subj.I-1* | 0 | 1 | 0 | 1 | 0 | 0 |
| *Subj.I* | 0 | 0 | 1 | 0 | 0 | 1 |

*Note.* Illustrative example of nominal (a and c) and disjunctive (b and d) coding of illustrative SNPs and diagnosis (Dx) and APOE. For SNP 1, all genotypes have a sufficient frequency and are coded (à la genotypic model), but for SNP 2, the minor homozygote (CC) does not occur frequently enough and is thus combined with the heterozygote (à la dominant model). In all tables, a 1 indicates the presence of a particular level of a categorical variable while a 0 indicates absence (e.g., *Subj.1* is an Alzheimer's Disease patient, with 2 ApoE E4 alleles, the AG genotype for SNP1 and either a CC or CA [presence of minor allele] for SNP2). Note that one subject has missing data (i.e., "<NA>"). This subject's data for SNP 1 is imputed to the mean of the sample for that SNP (SNP 1) where AA occurs in 70% of the sample, AG in 20%, and GG in 10%, therefore the missing data are imputed to those values.

We used two forms of PLS-CA in our discovery phase: discriminant PLS-CA and seed PLS-CA. We used discriminant PLS-CA in the validation phase. For details on background, and notation on PLS-CA and its derivatives see [31]. We briefly describe our motivation to use these two techniques here. We expand on the motivation in *Study design and overview*. Discriminant PLS-CA is a technique that maximizes the separation between *a priori* groups of participants (see also [35]). We used Discriminant PLS-CA to identify genotypes most associated with each group. In Seed PLS-CA, a “seed” is a specific genetic marker, where the seed analysis looks for distributions of genotypes similar to the seed (i.e., linkage disequilibrium). We used seed PLS-CA to identify other genotypes with distributions similar to *APOE*, which is the strongest contributor to non-familial AD [36,37]; thus we were trying to find additional candidate genotypes that have roughly the same pattern as *APOE* in the ADNI sample.

### 2.3 SNP Quality Control & Preprocessing

For all analyses we excluded any SNPs in the X and Y chromosomes, the pseudoautosomal region (XY), and mitochondrial region (i.e., we analyzed SNPs in Chromosomes 1-22). All SNPs were preprocessed with PLINK (v1.07; [38]) and in-house R code. SNP annotation was performed with the NCBI2R [39], and biomaRt packages [40,41]. We used both because in some cases, one annotation package would have information the other did not.

Participant and SNP call rates (i.e., completeness of data) ≥ 90%, minor allele frequency ≥ 5%, Hardy-Weinberg equilibrium *p ≤* 10^-6^. SNPs were then recoded into a disjunctive format (see Table 2a and b). Additionally, any genotype below a 5% threshold was combined with another genotype. In our study here, only the minor homozygotes were below the 5% threshold, and thus combined with the heterozygotes, which is analogous to a dominant model. Missing genotypes were imputed to the mean of the sample (see Table 2a and b).

### 2.4 Study design and overview

We conducted a two-part study: *Discovery* and *Validation*. In the *Discovery* phase there were two analyses with ADNI-1 genome-wide SNPs. The results from the *Discovery* analyses in ADNI-1 were used to create candidate SNP panels for validation in ADNI-GO/2. In the validation phase there was one analysis with a specific subset of ADNI-GO/2 SNPs.

Data from ADNI-1 and ADNI-GO/2 were not combined or preprocessed together at any stage in this study, so that no contamination or influence occurred from one set on the other. The two samples were also kept separate in order to preserve statistical independence for the discovery-validation pipeline. Because ADNI-1 and ADNI-GO/2 have two different chip sets, we generated a candidate panel of SNPs for ADNI-GO/2 based on the SNPs and their associated genes identified in ADNI-1 *(Discovery)*.

#### 2.4.1 Discovery study analyses

The goal of the diagnosis (Dx) × genotype analyses was to detect genotypes most associated with each diagnostic category. The seed PLS-CA used *APOE* E4 as the seed, and was performed on *APOE* (0, 1, or 2 E4 alleles) × genotypes. Both analyses were designed to identify candidate markers of AD: the discriminant analysis (henceforth referred to as “Dx-GWAS”) identifies genotypes most associated with each group, whereas the seed analysis (henceforth referred to as “ApoE-GWAS”) identifies genotypes similar to *APOE*. All analyses used bootstrap resampling [42] to identify stable genotypes. The distributions around the genotype were tested with “bootstrap ratios” (BSR; [31]). Significant genotypes in our two GWAS (in ADNI-1) were then used to create a new candidate panel of SNPs for validation (in ADNI-GO/2).

#### 2.4.2 Creation of SNP panels from Discovery for Validation

Because ADNI-1 and ADNI-GO/2 were used as independent data sets in our study, and because the data come from two different genome-wide chips, we used the significant genotypes from the discovery analyses (i.e., Dx-GWAS and the ApoE-GWAS) to generate candidate SNPs for validation. For all significant genotypes in the discovery analyses, we used their respective SNPs to: (1) compile a list of all SNPs within a 50kbase (25+/-) window of those SNPs, and (2) retrieve all stable ensembl gene (ENSG) IDs associated with those SNPs from the discovery analyses, and in turn retrieve all possible SNPs associated those ENSG IDs. All SNPs from the steps (1) and (2) were combined into a candidate set. We extracted all SNPs from the ADNI-GO/2 data that were from the discovery-derived candidate set for use in the validation phase.

#### 2.4.3 Validation study analyses

Validation analyses were conducted on the ADNI-GO/2 data set with a discriminant PLS-CA on the validation SNPs based on diagnosis. The goal of the validation Dx × genotype analysis was to detect genotypes most associated with each diagnostic category. As in the discovery phase, we used bootstrap resampling and the BSR test to identify stable genotypes.

## 3. Results

### 3.1 Discovery (ADNI-1)

ADNI-1 genome-wide data contains 620,901 SNPs and 757 participants. After QC and preprocessing 756 participants (AD = 344, MCI = 204, CON = 208) and 517,441 SNPs (in chromosomes 1-22) remained, which produced a 756 participants × 1,332,455 disjunctive genotypes matrix (see Table 2). Only the first two MCA components showed stratification effects (race and ethnicity) and were thus removed from (i.e., regressed out of) the data. Subsequent components showed no apparent effects of stratification. For the discovery GWAS, we used a cutoff of ±5 for the BSR tests which is slightly below the traditional GWAS parametric threshold *(p*< .05 × 10^-8^ would correspond to a BSR ≈ 5.33).

#### 3.1.1 Dx-GWAS

The discriminant PLS-CA produced two components. Component 1 explained 50.25% of the variance and was driven by the separation of the AD group from the MCI group (see Supplemental Figure 1). Component 2 explained 49.76% of the variance and separated the CON group from the other two groups (see Supplemental Figure 1). Figure 1 shows all genotypes plotted with their BSR values in a Manhattan-like plot. BSR values can be positive or negative— because the sign matches their component score—we call this plot “Manhattan on the Hudson” (MotH; like a city skyline and reflection on a river). Figure 1a shows the BSRs for all genotypes for Dx-GWAS Component 1 and Figure 1b shows the BSRs for all genotypes for Dx-GWAS Component 2. The majority of stable genotypes were more related to the AD group than the other groups (see Figure S1 and Table S1).

**Figure 1.**
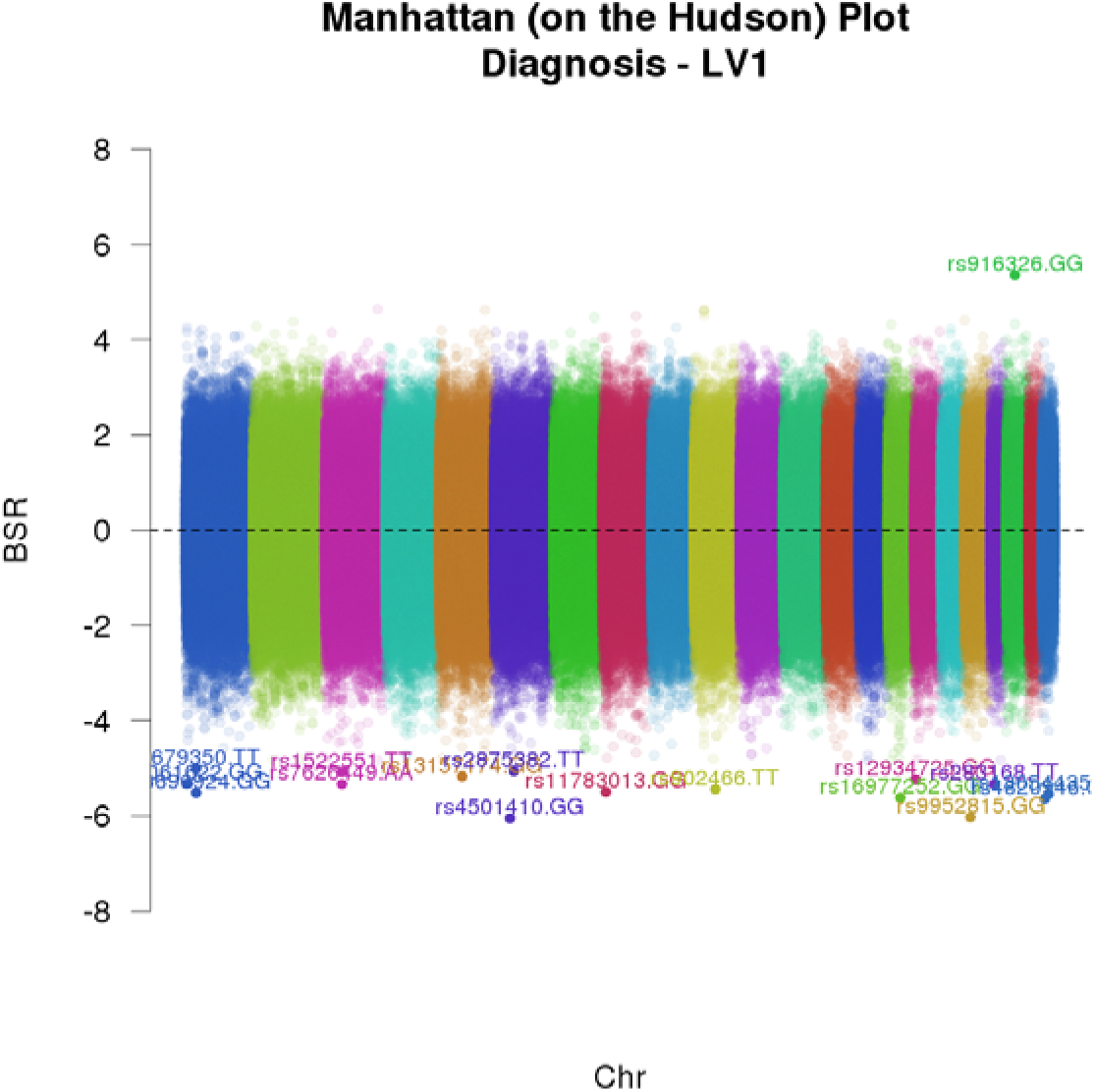
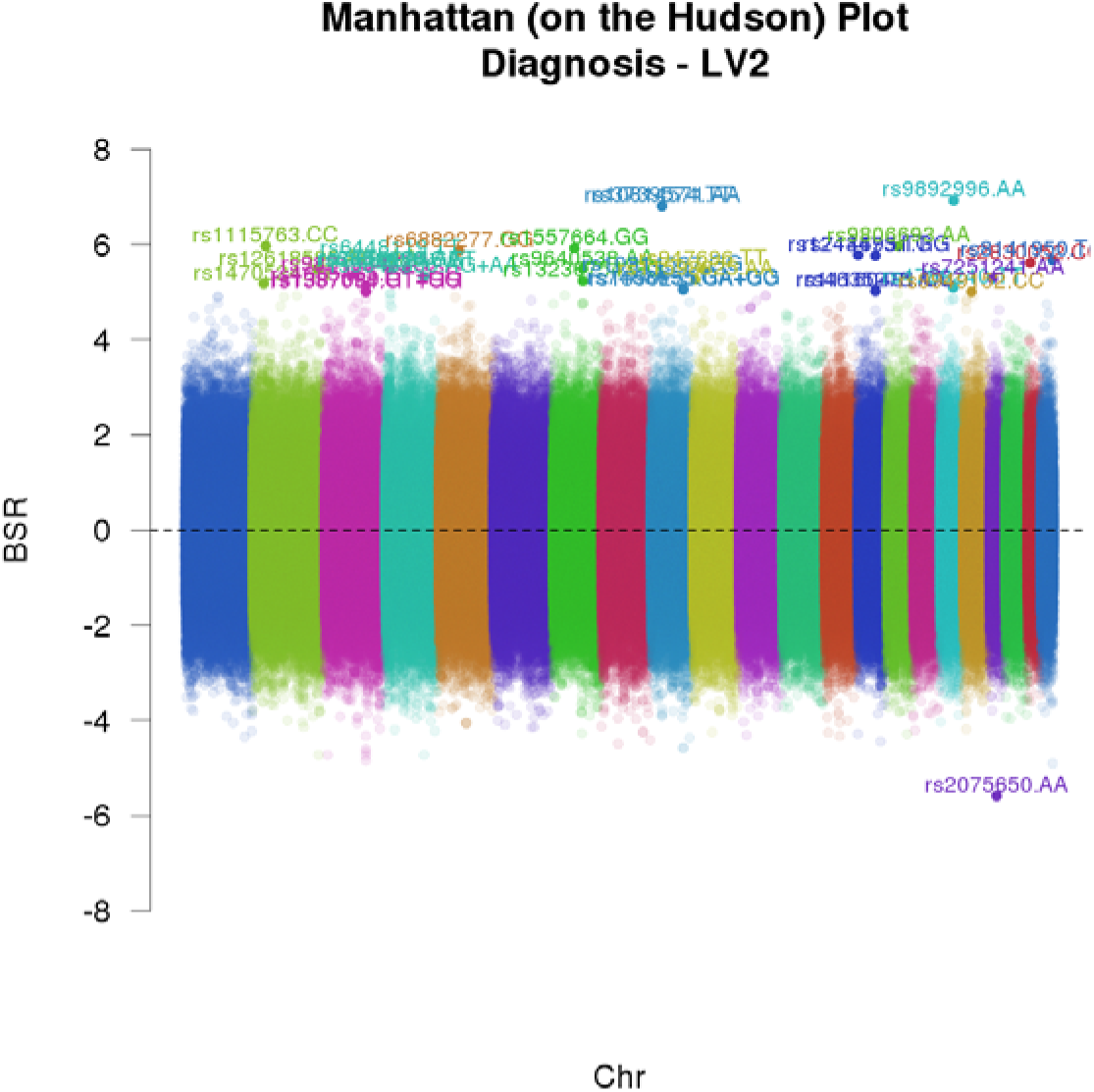
Manhattan (on the Hudson) plots for the multivariate Diagnosis (Dx)-GWAS. Horizontal axes are each genotype ordered by Chromosome (Chr), where each Chr is color-coded (1-22). The vertical axes are bootstrap ratio values (BSRs). Both panels show BSRs (analogous to *t*- or *Z*-scores; which can be positive or negative) for each genotype along Component 1 (a.k.a. Latent Variable (LV) 1; panel a, top) and Component 2 (a.k.a. LV 2; panel b, bottom) - the same components as in Figure 2. With respect to the multivariate Dx-GWAS a wide variety of genotypes show significant, and similar, effects and are not concentrated in any particular region (see also Table S1 and Figure S1).

#### 3.1.2 ApoE-GWAS

Because there were only 3 levels to the seed (“0 E4,” “1 E4,” and “2 E4” alleles), seed PLS-CA produced only two components. Component 1 explained 51.24% of the total variance and was driven by the presence (left) vs. absence (right) of E4 alleles (see Supplemental Figure 2). Component 2 explained 48.76% of the variance and separates the two E4 alleles group from the other two groups (see Supplemental Figure 2). Figure 2 shows the BSRs for all genotypes from this analysis in a MotH plot. Figure 2a shows the BSRs for all genotypes for ApoE-GWAS Component 1 and Figure 2b shows the BSRs for all genotypes for ApoE-GWAS Component 2 (see also Figure S2 and Table S2).

**Figure 2.**
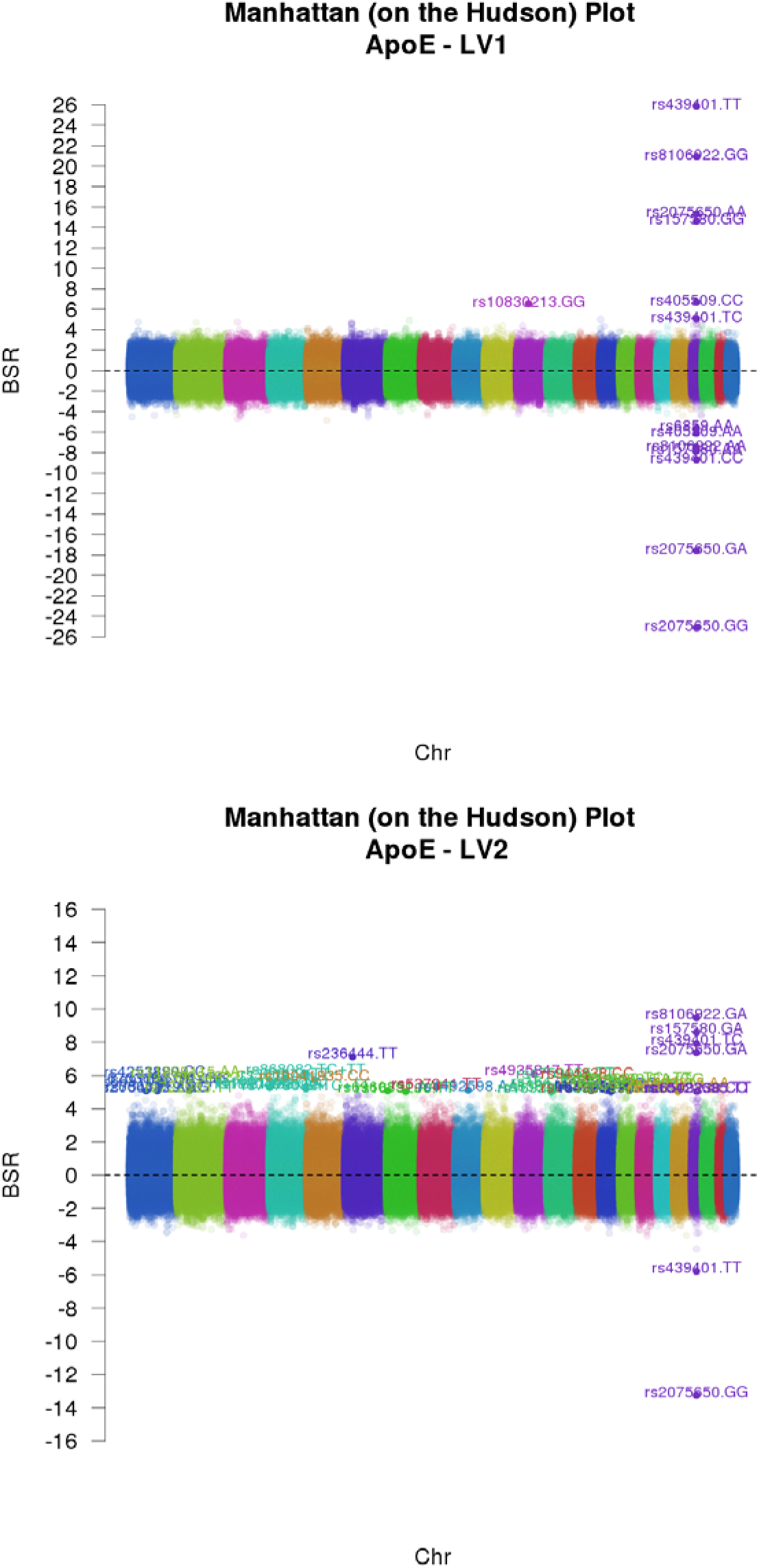
Manhattan (on the Hudson) plots for the multivariate ApoE-GWAS. Horizontal axes are each genotype ordered by Chromosome (Chr), where each Chr is color-coded (1-22). The vertical axis is bootstrap ratio values (BSRs). Both panels show BSRs (analogous to *t*- or *Z*-scores; which can be positive or negative) for each genotype along Component 1 (a.k.a. Latent Variable (LV) 1; panel a, left) and Component 2 (a.k.a. LV 2; panel b, right) - the same components as in Figure 4. With respect to the multivariate ApoE-GWAS, many of the effects are concentrated, generally, in Chr19 (see also Table S2) across both components, but much more so for Component 1 (a; left). While Component 2 (b; right) shows a variety of effects, some of the strongest are still in Chr19.

### 3.2 Candidate panel creation

All SNPs associated with significant genotypes in *Discovery* were used to create the candidate panel. A total of 105 genotypes from 96 SNPs exceeded the ±5 BSR threshold (see Supplemental Tables 1 and 2). From these 96 SNPs, our candidate panel process identified 1,045,360 possible SNPs.

### 3.3 Validation (ADNI-GO/2)

ADNI-GO/2 genome-wide data contains 730,525 SNPs and 791 participants. We extracted 5,508 SNPs from the ADNI-GO/2 chipset based on the candidate panel of 1,045,360 SNPs. After QC and preprocessing, 791 participants (AD = 203, MCI = 319, CON = 269) and 5,508 SNPs remained, which produced a 791 participants × 14,200 disjunctive genotypes matrix (see Table 2). Only the first two MCA components showed stratification effects (race and ethnicity) and were thus removed from the data. Subsequent components showed no apparent effects of stratification.

A discriminant PLS-CA (Dx × genotypes) was performed on the 791 × 14,200 matrix. For the validation analysis, we used a cutoff of ±3.25 for the BSR tests (roughly equivalent to a Bonferroni cutoff for the number of unique genes). Discriminant PLS-CA produced two components: Component 1 explained 50.41% and was driven by the separation of the MCI group from the AD and CON groups (see Figure 3); Component 2 explained 49.59% of the variance and separated the CON group from the AD group (see Figure 3).

**Figure 3.**
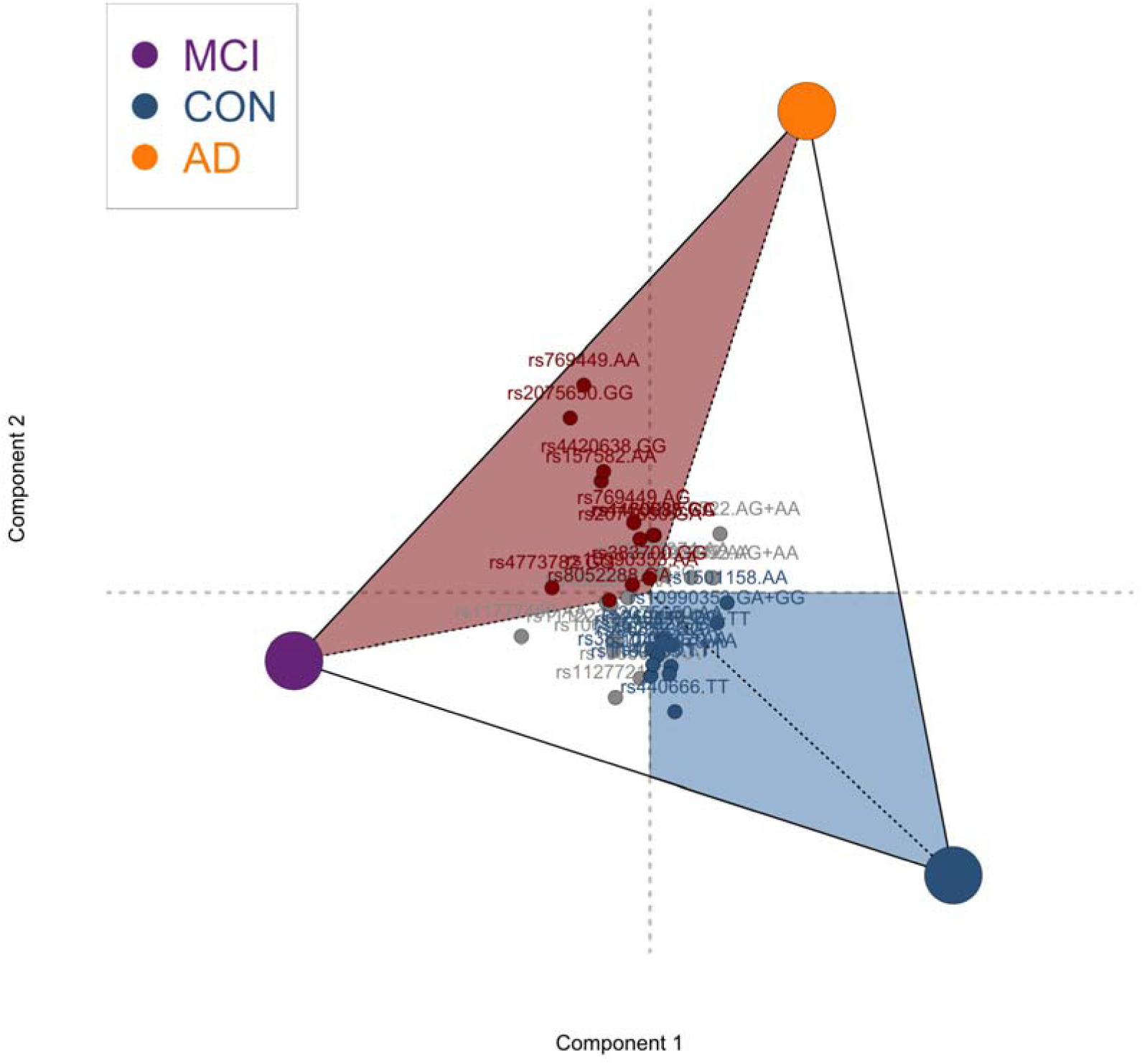
Diagnosis-based analysis in the *Validation* phase. The groups form a boundary region on the components. We denoted portions of this subspace as “control” associated (blue) or “risk” associated (red); anything outside the control or risk regions is ambiguous. Genotypes that fall in the “control” region are more associated with the control group than the other groups and genotypes that fall in the “risk” region are more associated with {MCI or AD} than the CON group.

**Table 3.** *Validation set: significant genotypes* on either component

| rsid | genotype | BSR - 1 | BSR - 2 | chr | Gene symbol | Risk / Control Region |
| --- | --- | --- | --- | --- | --- | --- |
| rs10494979 | AA | 0.95 | -3.422 | 1 | PTPN14 | CON |
| rs11122374 | AA | 1.985 | 3.255 | 1 | TSNAX-DISC1,DISC1 |  |
| rs11122374 | GA+GG | -1.995 | -3.346 | 1 | TSNAX-DISC1,DISC1 |  |
| rs1501158 | AA | 3.319 | -0.409 | 4 | LINC00504 | CON |
| rs1065261 | TT | -1.971 | -3.451 | 6 | CPNE5 |  |
| rs3213537 | TC+TT | 1.345 | -3.31 | 6 | CPNE5 | CON |
| rs11777456 | AA | -3.366 | -1.615 | 8 | NCALD |  |
| rs10990353 | AA | -3.374 | 1.483 | 9 | ZYG11AP1 / LOC100421294 | RISK |
| rs10990353 | GA+GG | 3.496 | -1.482 | 9 | ZYG11AP1 / LOC100421294 | CON |
| rs7093342 | GA | -3.395 | -0.848 | 10 | ITIH5 |  |
| rs7093342 | AA | 3.904 | 1.5 | 10 | ITIH5 |  |
| rs1979522 | AG+AA | 3.275 | 2.298 | 12 | LRMP |  |
| rs4773782 | GG | -3.608 | 0.185 | 13 | GPC6 | RISK |
| rs17105992 | GG | -3.513 | -0.847 | 14 | LOC107984016,RAD51B |  |
| rs17105992 | AG+AA | 3.55 | 0.844 | 14 | LOC107984016,RAD51B |  |
| rs4902611 | AA | -3.93 | -1.293 | 14 | LOC107984016,RAD51B |  |
| rs8052288 | GA | -3.476 | -0.692 | 16 | RBFOX1 | RISK |
| rs1553614 | AA | -0.278 | -3.272 | 16 | RBFOX1 |  |
| rs6859 | GG | 0.415 | -3.895 | 19 | NECTIN2‡ | CON |
| rs2075650 | GA | -0.656 | 3.5 | 19 | TOMM40 | RISK |
| rs2075650 | AA | 1.555 | -5.558 | 19 | TOMM40 | CON |
| rs2075650 | GG | -2.12 | 4.338 | 19 | TOMM40 | RISK |
| rs157582 | GG | 0.162 | -5.297 | 19 | TOMM40 | CON |
| rs157582 | AA | -1.773 | 3.72 | 19 | TOMM40 | RISK |
| rs1160985 | CC | 0.344 | 4.349 | 19 | TOMM40 | RISK |
| rs1160985 | TT | 0.048 | -4.101 | 19 | TOMM40 | CON |
| rs769449 | GG | 1.749 | -6.527 | 19 | APOE | CON |
| rs769449 | AA | -1.792 | 5.195 | 19 | APOE | RISK |
| rs769449 | AG | -0.995 | 4.194 | 19 | APOE | RISK |
| rs439401 | TT | 0.678 | -3.277 | 19 | APOE | CON |
| rs4420638 | GA | 0.207 | 3.95 | 19 | APOC1 | RISK |
| rs4420638 | AA | 0.803 | -6.458 | 19 | APOC1 | CON |
| rs4420638 | GG | -1.637 | 3.913 | 19 | APOC1 | RISK |
| rs383700 | GG | -0.179 | 3.314 | 21 | APP | RISK |
| rs383700 | AG+AA | 0.179 | -3.436 | 21 | APP | CON |
| rs440666 | TT | 0.631 | -3.88 | 21 | APP | CON |
| rs1127721 | TT | -0.789 | -3.543 | 22 | PARVG |  |

*Note.* ‡ rs6859 previously associated with PVRL2. Significant genotypes for both components from the validation set. Component 1 primarily separated CON from AD, whereas Component 2 primarily separated AD from the other two groups. In the validation analysis, we used the asymmetric version of correspondence analysis (CA) to help visualize which genotypes are most associated with each group, and to create identify “control” (CON) and “risk” (RISK) regions within the CA components (see Figure 3). Genotypes labeled as CON or RISK indicate which region they are in, otherwise their association is more ambiguous in terms of association with risk or control (“protective”) factors.

Table 3 lists all significant genotypes from *Validation*. In Figure 3, we highlight boundaries to interpret genotypic effects. We focused specifically on two boundaries: (1) the boundary associated with the CON group (lower right, blue) and (2) the boundary associated with “risk status” (i.e., AD or MCI; upper middle to middle left, red). The relevant SNPs are highlighted in their boundary colors in Figure 3 and identified as part of the “RISK” or “CON” region in Table 3.

## 4. Discussion

There has been an increased interest in multivariate approaches for genetics [43], especially for AD [44]. However, many studies in AD still only use the additive model contrary to: (1) known non-linear and non-additive effects (such as in [45–50]), and (2) the fact that the genotypic [19] or co-dominant [20,51] models appear to be better suited for complex effects.

Because of our approach we easily identified more specific genetic contributions to AD than other approaches can. We emphasized genotypes instead of presumed additive effects within SNPs. Our *Validation* phase revealed many complex effects that highlight why a simple additive model (all in the same presumed direction, i.e., the minor allele), or any *a priori* model, may not be sufficient to characterize complex genetic effects (see also Figure 3 and Table 3); for examples: (1) rs769449 is approximately linear where the minor homozygote and the heterozygote are in the risk region, and the major homozygote in the control region, (2) rs4420638 is approximately dominant where the minor allele is in the risk region, (3) rs440666 is approximately recessive where the minor homozygote is in the control region, and (4) rs8052288 showed a heterozygous effect in the risk region.

### 4.1 Specific effects of well-known genes

Some of the strongest effects in all of our analyses were associated with genotypes in Chromosome 19 (Chr19). These effects included some well-known genes in AD: *TOMM40, APOE, APOC*, and *NECTIN2*. However, PLS-CA identified exactly which genotypes contributed to which effects (see Fig. 3 and Table 3). For examples, the ‘AA’ genotype of rs769449 *(APOE)* and the ‘GG’ genotype of rs2075650 *(TOMM40)* are at the extreme of our plot, directly opposite of the CON group, and almost exactly half way between AD and MCI (Fig. 3). This means that ‘AA’ in rs769449 and GG in rs2075650 rarely occur in CON, but tend to occur roughly equally in both the AD and MCI groups. Most importantly for Chr19 effects: our *Discovery* phase identified ‘AA’ of rs6859 associated with the *presence* of *APOE* E4, whereas our *Validation* phase identified ‘GG’ of rs6859 associated with the CON group. While the “direction” of the effect is the same across both studies, the source of the effect was not (i.e., minor allele vs. major allele). Taken together, the *Discovery* and *Validation* analyses suggest that, depending on the genotype, rs6859 confers both a risk and a possibly protective factor.

*APP* is also well-known in AD [52]. We found that *APP* was identified through the Dx-GWAS in *Discovery* and also showed effects in the *Validation* analyses (see Table 3). The *Validation* analyses show that various *APP* genotypes were associated with “risk” and “control” (Fig. 3 and Table 3). Given the findings across both our *Discovery* and *Validation* phases, our findings here suggest that *APP* is more related to diagnostic criteria than to *APOE* distribution, and that specific *APP* genotypes provide possibly protective effects (e.g., ‘TT’ in rs440666).

### 4.2 Lesser-known and novel genetic effects

We found risk specific genotypes from SNPs in the *GPC6* and *RBFOX1* genes (Table 3; Fig. 3). Both of these genes have been associated with pathological or cognitive phenotypes of AD. These two genes have shown associations with very different phenotypic or outcome measures in AD. *RBFOX1* has been associated with pathological effects in the brain, such as neurofibrillary tangles [53] as well as neuroimaging phenotypes [54] and hippocampal volumes [55] in AD. In contrast, *GPC6* has been associated with cognitive and behavioral decline in AD [56].

Our analyses also revealed several novel genetic effects. Some of these were “control” effects, and some were “risk” effects. Of the control-associated effects, we found contributions from genotypes in the *PTPN14, CPNE5*, and *LINC00504* genes. Additionally, we also found contributions from genotypes in the *ZYG11AP1* and *LOC100421294* genes, where each had various genotypes associated with both “control” and “risk” effects. There are no reported effects of rs1501158 *(LINC00504)* and rs10990353 *(ZYG11AP1* / *LOC100421294)* in AD, dementia, or the cognitive aging literature. *CPNE5* has not been reported in AD but has been associated with other neurodegenerative diseases [57].

Finally, there were also several stable effects that were “ambiguous” (i.e., we could not classify as “risk” or “control” because they fall outside of our designated regions). However, these effects are of interest because (1) effects appear in both analyses and (2) of their associations in various neurodegenerative disorders or their interactions with other genes that play substantial roles in AD. *PARVG* has been associated with Parkinson’s Disease [58] and neurodegeneration [59]. Furthermore, *RAD51B, NCALD*, and *DISC1* are rarely reported in AD, but they have been associated with the amyloid precursor gene *APP* [60]. *RAD51B* has also been associated with macular degeneration [61,62]. Furthermore, *DISC1* has been associated with late-onset AD [63], as well as Aβ production [64].

## 5. Conclusions

Like in our work here, [54–56] use the ADNI sample. However, these studies use more complex study designs or more complex methods, which, in some cases [54,55], demand vast computational resources. Other work, such as [56], also have more data and larger samples. Furthermore, [65] also used the ADNI sample and have recognized the utility and power of the genotypic model. Yet, compared to all of these approaches, our much simpler and computationally less expensive approach (with a comparable or smaller sample size than these other studies) found some of the same effects reported in all of the aforementioned papers, such as the contributions of *GPC6* and *RBFOX1*. PLS-CA also found a number of well-known genetic contributions to AD such as the effects across Chr19. The Chr19 effects were important because they highlight that our method can and does identify well-known and robust effects, and thus illustrates validity for the problems at hand. But more importantly, we identified many novel effects that otherwise could not be detected without a more general approach. Taken together, our study helps reveal some of the genetic complexities of AD, especially in that we have identified many genotypes from many genes that contribute in a variety of ways (e.g., minor allele as possibly protective, heterozygous effects).

PLS-CA has many advantages over traditional and more recent approaches to GWAS. First, PLS-CA does not make assumptions about genotypic effects, rather, PLS-CA reveals the types of effects (e.g., additive, dominant) and the directions of these effects. More importantly, because PLS-CA is a multivariate technique, it actually provides estimates of the contributions of these genotypes to polygenic effects (see BSRs in Table 3 or component scores in Figure 3). Together these features of PLS-CA are particularly suited for our problem. PLS-CA has identified and specified the expected complex genetic contributions to AD, and thus techniques such as PLS-CA help making clearer interpretations, as well as reduce false positives, non-replications, conflicting reports, and otherwise problematic interpretations of genetic effects.

### 5.1 Limitations

The ADNI data sets are, by today’s standards, relatively small samples for such a study. However, our study design emphasized our *Validation* step to help confirm effects identified in *Discovery*. Additionally, our *Validation* phase could have weighted the candidates based on bootstrap ratios from the *Discovery* phase to emphasize the strength of effects. However, we chose not to and opted for a more data-driven strategy. Ultimately this choice was beneficial: for example, if we had used weighted values in the *Validation* phase we may have missed effects such as those associated with rs6859 (i.e., rs6859 expressed different effects in *Discovery* compared to *Validation)*. Furthermore, the individuals within these groups are heterogeneous, and could possibly have confounding factors (e.g., vascular incidents) or misdiagnoses. Because of the difficulty of diagnosing AD *in vivo*, we are limited in our claims to AD broadly, but have provided a clearer genetic landscape of the cohorts within ADNI-1 and ADNI-GO/2. Thus, the effects we have identified should be further verified in a predictive analysis wherein individuals could be genotyped for the specific markers we have identified; which may be possible through similar studies that are in progress (e.g., ADNI-3, the Ontario Neurodegenerative Disease Research Initiative [66]).

## Acknowledgements

Data collection and sharing for this project was funded by the ADNI (National Institutes of Health Grant U01 AG024904) and DOD ADNI (Department of Defense award number W81XWH-12–2–0012). ADNI is funded by the National Institute on Aging, the National Institute of Biomedical Imaging and Bioengineering, and through generous contributions from the following: Alzheimer’s Association; Alzheimer’s Drug Discovery Foundation; Araclon Biotech; BioClinica, Inc.; Biogen Idec Inc.; Bristol-Myers Squibb Company; Eisai Inc.; Elan Pharmaceuticals, Inc.; Eli Lilly and Company; EuroImmun; F. Hoffmann-La Roche Ltd and its affiliated company Genentech, Inc.; Fujirebio; GE Healthcare; IXICO Ltd.; Janssen Alzheimer Immunotherapy Research & Development, LLC.; Johnson & Johnson Pharmaceutical Research & Development LLC.; Medpace, Inc.; Merck & Co., Inc.; Meso Scale Diagnostics, LLC.; NeuroRx Research; Neurotrack Technologies; Novartis Pharmaceuticals Corporation; Pfizer Inc.; Piramal Imaging; Servier; Synarc Inc.; and Takeda Pharmaceutical Company. The Canadian Institutes of Health Research is providing funds to support ADNI clinical sites in Canada. Private sector contributions are facilitated by the Foundation for the National Institutes of Health (www.fnih.org). The grantee organization is the Northern California Institute for Research and Education, and the study is coordinated by the Alzheimer’s Disease Cooperative Study at the University of California, San Diego. ADNI data are disseminated by the Laboratory for Neuro Imaging at the University of Southern California. For up-to-date information and data see: http://www.adni-info.org and http://adni.loni.usc.edu/

## Supplemental Material

**Table S1.**
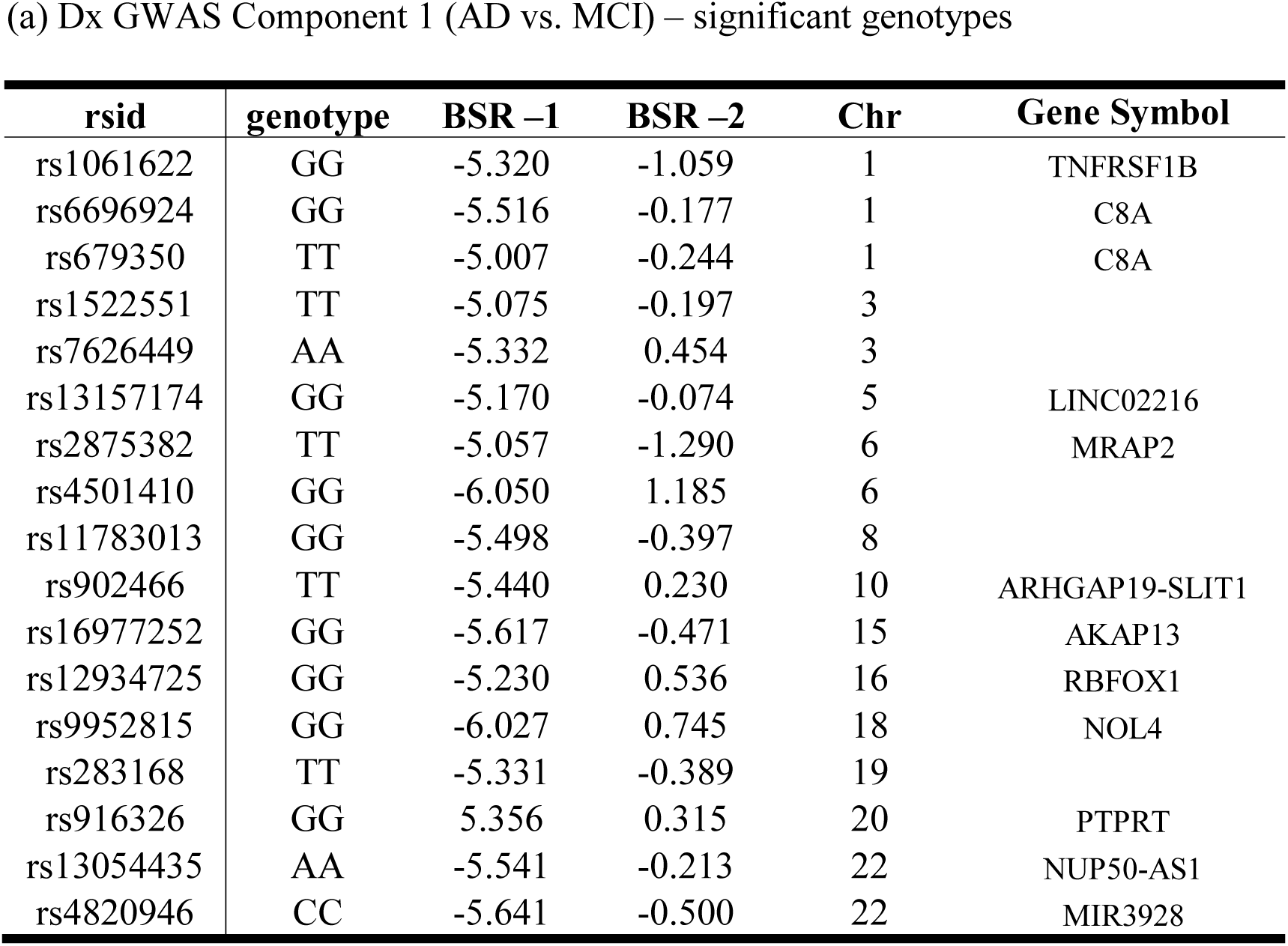
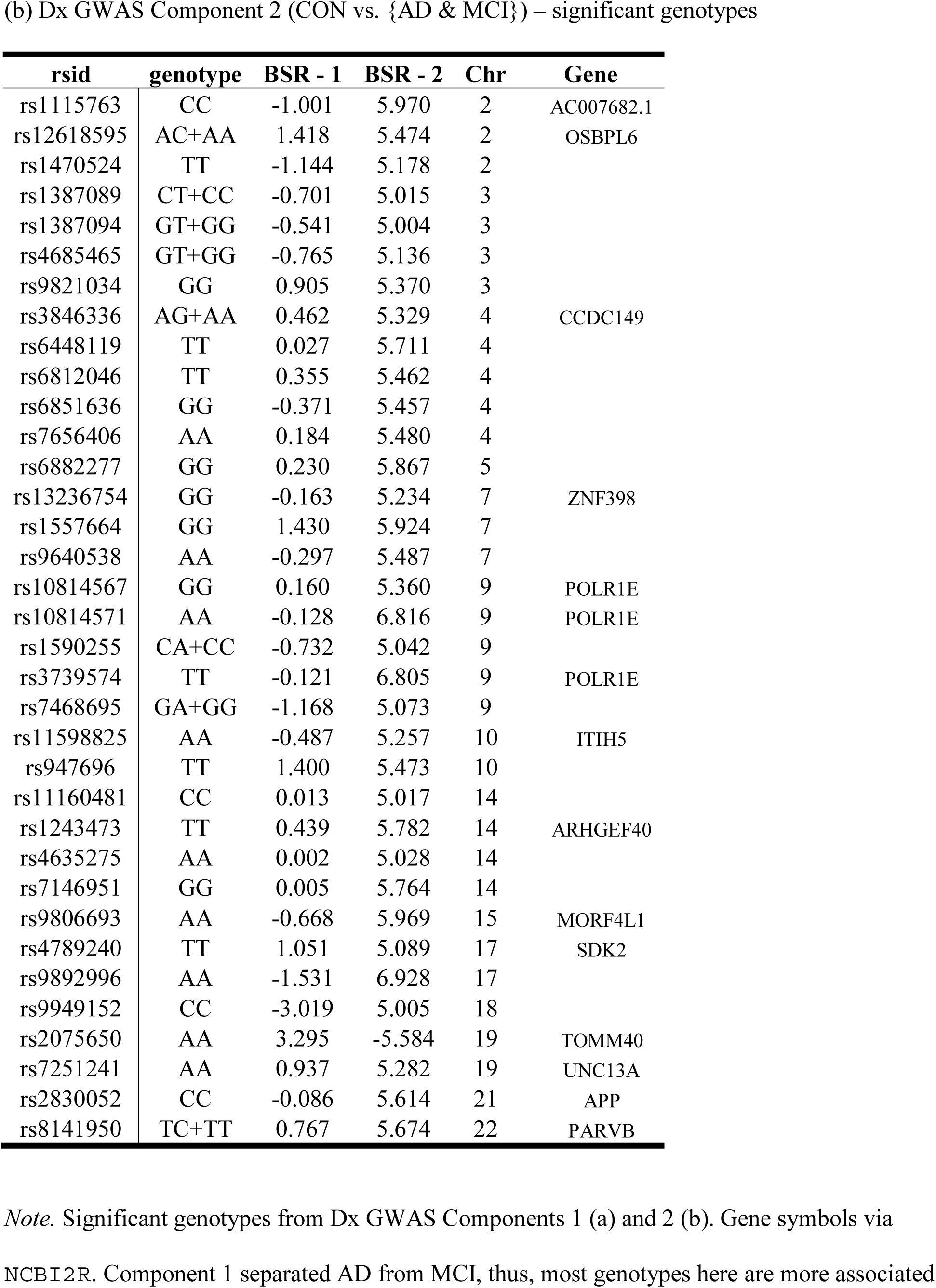
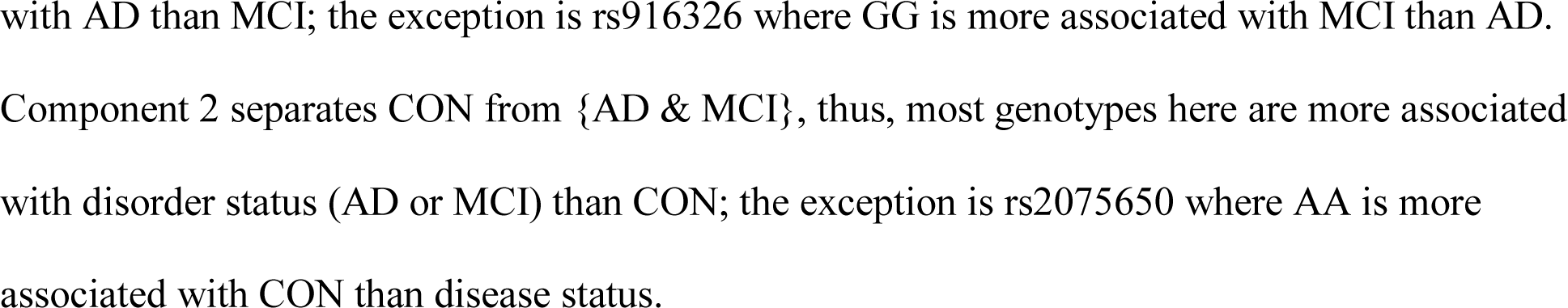
Dx GWAS Significant genotypes

**Table S2.** ApoE GWAS Significant genotypes

| **rsid** | **genotype** | **BSR - 1** | **BSR - 2** | **Chr** | **Gene Symbol** |
| --- | --- | --- | --- | --- | --- |
| rs10830213 | GG | 6.537 | -0.724 | 11 | RAB38 |
| rs157580 | AA | -7.842 | -2.918 | 19 | TOMM40 |
| rs157580 | GG | 14.612 | -2.130 | 19 | TOMM40 |
| rs2075650 | AA | 15.323 | -0.619 | 19 | TOMM40 |
| rs405509 | AA | -6.101 | -2.567 | 19 | APOE |
| rs405509 | CC | 6.701 | -1.244 | 19 | APOE |
| rs439401 | CC | -8.703 | -2.630 | 19 |  |
| rs6859 | AA | -5.614 | -3.096 | 19 | NECTIN2/PVRL2 |
| rs8106922 | AA | -7.514 | -3.490 | 19 | TOMM40 |
| rs8106922 | GG | 20.928 | -4.430 | 19 | TOMM40 |

(b) ApoE GWAS Component 2 (2 E4 alleles vs. {0 or 1}) – significant genotypes
| rsid | genotype | BSR - 1 | BSR - 2 | Chr | Gene Symbol |
| --- | --- | --- | --- | --- | --- |
| rs2000072 | AA | -0.211 | 5.066 | 1 | LINC00624 |
| rs4253890 | CC | -1.077 | 5.985 | 1 | PTPN14 |
| rs6681032 | TC+TT | 0.157 | 5.439 | 1 |  |
| rs6703696 | GA+GG | 0.169 | 5.604 | 1 |  |
| rs7541019 | GG | 0.576 | 5.069 | 1 | TSNAX-DISC1,DISC1 |
| rs11899115 | AA | -0.640 | 5.996 | 2 |  |
| rs13009482 | CT+CC | -1.383 | 5.696 | 2 |  |
| rs722963 | TT | -0.288 | 5.093 | 2 |  |
| rs10019637 | CC | -0.964 | 5.412 | 4 |  |
| rs10804966 | AA | -0.151 | 5.267 | 4 | EVC |
| rs300574 | TT | 0.275 | 5.860 | 4 | SPRY1 |
| rs7678082 | TC+TT | -0.105 | 5.196 | 4 | WWC2 |
| rs7681283 | GG | -0.168 | 5.278 | 4 | EVC |
| rs868082 | TC+TT | -0.187 | 6.130 | 4 |  |
| rs10041935 | CC | -0.895 | 5.750 | 5 |  |
| rs236444 | TT | -1.938 | 7.117 | 6 | CPNE5 |
| rs1673206 | TT | -1.294 | 5.026 | 7 |  |
| rs6960851 | AA | -0.713 | 5.079 | 7 |  |
| rs537941 | TT | -0.730 | 5.270 | 8 | NCALD |
| rs1492598 | AA | -0.933 | 5.119 | 9 |  |
| rs4935847 | TT | -0.247 | 6.085 | 11 |  |
| rs1647147 | GG | -1.396 | 5.407 | 12 |  |
| rs16928445 | TT | 0.609 | 5.015 | 12 | LRMP |
| rs4759955 | TT | -0.490 | 5.903 | 12 | TMEM132D |
| rs6582412 | AA | -1.433 | 5.113 | 12 |  |
| rs944838 | CC | 0.373 | 5.917 | 13 | GPC6 |
| rs9549831 | AA | 0.454 | 5.082 | 13 |  |
| rs4905290 | GG | -0.457 | 5.019 | 14 | CLMN |
| rs6573852 | GG | 0.438 | 5.294 | 14 | RAD51B |
| rs7148010 | TT | 1.018 | 5.112 | 14 | SMOC1 |
| rs10519492 | GA+GG | -0.468 | 5.537 | 15 |  |
| rs6496431 | GG | -0.087 | 5.072 | 15 |  |
| rs714900 | TC+TT | -0.627 | 5.611 | 15 |  |
| rs7177541 | AA | -0.664 | 5.299 | 15 |  |
| rs4500815 | AA | -0.280 | 5.300 | 18 | CTIF |
| rs9955327 | CC | -1.958 | 5.021 | 18 | CELF4 |
| rs10423685 | TT | -0.479 | 5.053 | 19 | ZNF600 |
| rs157580 | GA | 4.603 | 8.591 | 19 | TOMM40 |
| rs6509238 | CC | 0.056 | 5.038 | 19 |  |
| rs8106922 | GA | 3.703 | 9.456 | 19 | TOMM40 |

(c) ApoE GWAS Significant genotypes on both Components.
| rsid | genotype | BSR - 1 | BSR - 2 | Chr |
| --- | --- | --- | --- | --- |
| rs2075650 | GA | -17.593 | 7.374 | 19 |
| rs2075650 | GG | -25.132 | -13.27 | 19 |
| rs439401 | TC | 5.052 | 7.920 | 19 |
| rs439401 | TT | 25.848 | -5.790 | 19 |

*Note.* Significant genotypes from ApoE GWAS Components 1 (a), 2 (b), and on both Components (c). Gene symbols via NCBI2R. Component 1 separated presence from absence of E4 alleles. Genotypes that have a negative bootstrap ratio (BSR) were more associated with the presence of an E4 allele. Component 2 separated, essentially, the 2 E4 alleles from the other (0 or 1) E4 alleles. In (b) all genotypes were more related to the *absence* of 2 E4 alleles. In (c) these genotypes contribute to both components and suggest that these genotypes are in very high linkage disequilibrium with ApoE (note that the GG genotype of rs2075650 strongly contributes to both components and in the same direction as the *presence* 2 E4 alleles).

**Supplemental Figure 1.**
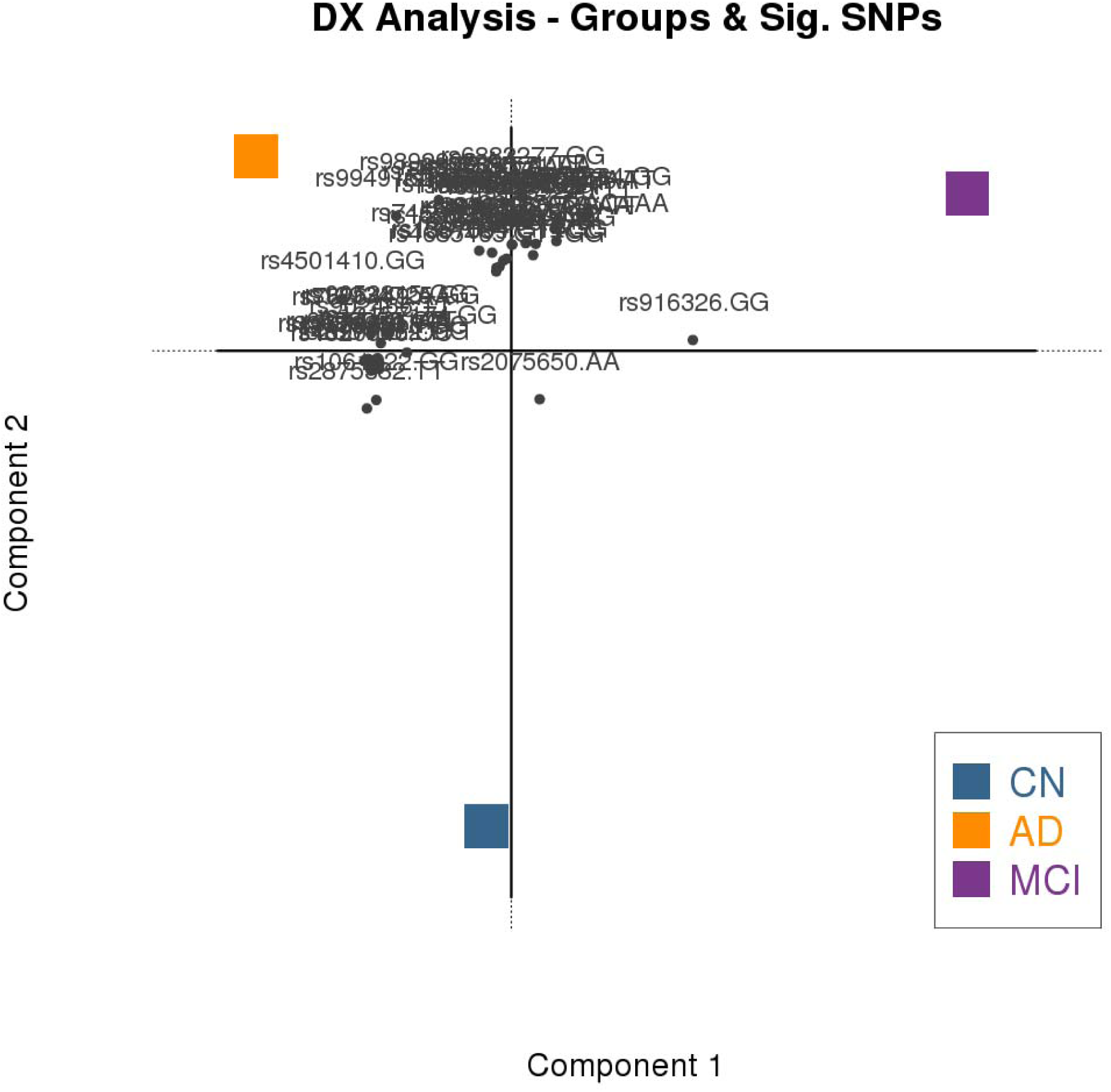
The component map from the Dx-GWAS (discovery phase). The component map shows significant genotypes and the group configuration to illustrate—as a biplot—the relationship between the genotypes and groups.

**Supplemental Figure 2.**
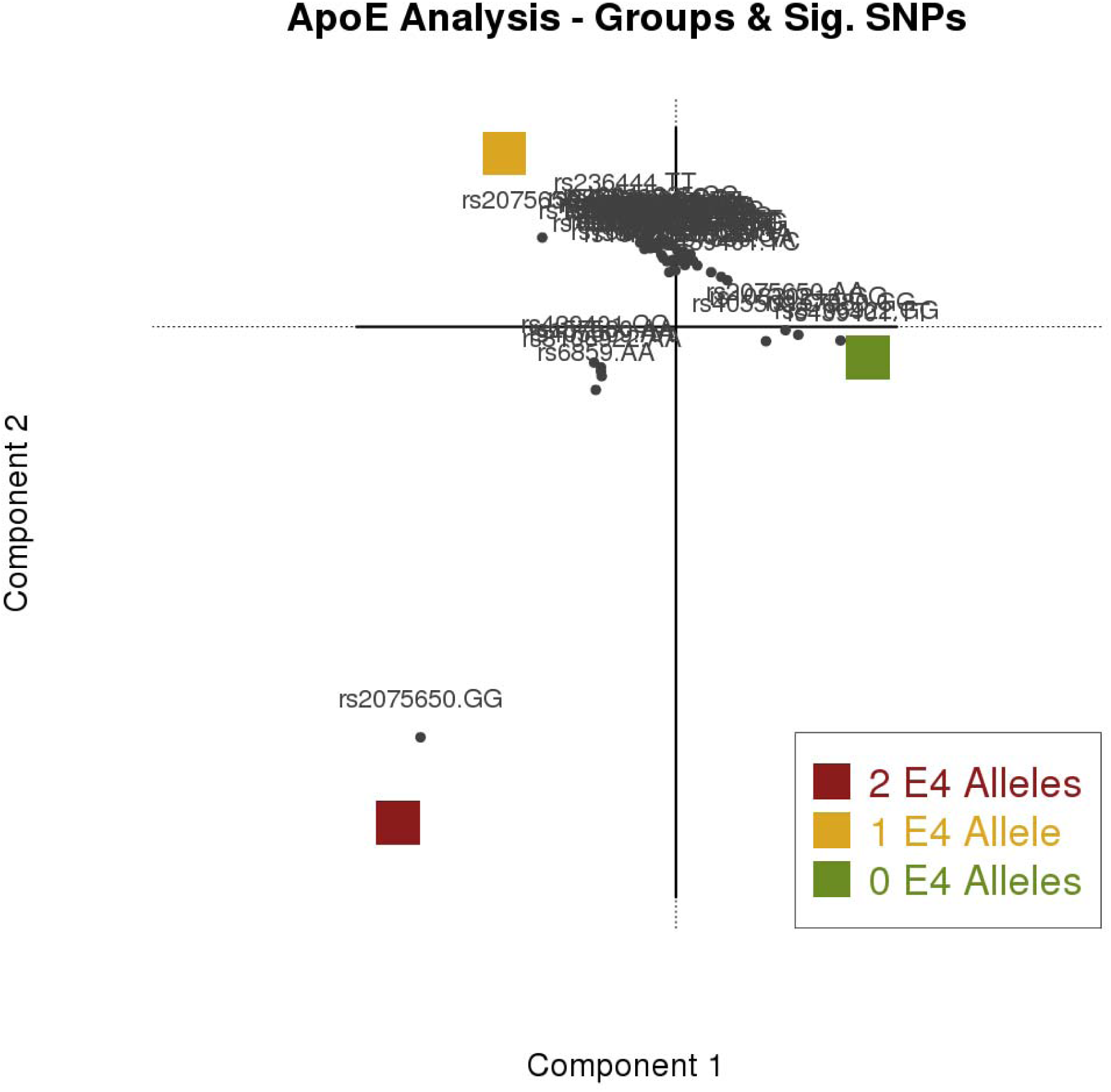
The component map from the ApoE-GWAS (discovery phase). The component map shows significant genotypes and the group configuration to illustrate—as a biplot—the relationship between the genotypes and groups.

